# Chronic activation of dopaminergic neurons via bioluminescence-optogenetics provides neuroprotection in a rodent model of Parkinson’s disease

**DOI:** 10.1101/2022.10.17.512538

**Authors:** Fu Hung Shiu, Henry Skelton, Ken Berglund, Alejandra M. Fernandez, Claire-Anne N. Gutekunst, Elizabeth R. Robinson, Zuhui Wang, Robert E. Gross

## Abstract

Previous studies in human patients and rodent models of Parkinson’s disease (PD) have established neuroprotection of dopaminergic (DA) neurons in substantia nigra pars compacta (SNC) by physical exercise, but the precise origin of this neuroprotective effect has yet to be elucidated. In this study, we tested a hypothesis that enhanced activity of DA neurons in SNC results in neuroprotection using the unilateral 6-hydroxydopamine (6-OHDA) injection model in mice. To increase activity of DA neurons chronically and specifically, we injected an adeno-associated viral vector carrying a step-function luminopsin (SFL) – a fusion protein of light-emitting *Gaussia* luciferase and light-sensing step-function channelrhodopsin 2 – into SNC ipsilateral to 6-OHDA using the pan-neuronal human synapsin I promoter or the Cre-lox system in transgenic mice expressing the recombinase under control of the tyrosine hydroxylase (TH) promoter. Upon application of SFL substrate, coelenterazine (CTZ), the luciferase moiety of luminopsin emits bioluminescence which in turn activate the opsin moiety. Daily injection of CTZ for 4 weeks ameliorated a stereotypical behavior, namely ipsiversive rotations, induced by unilateral 6-OHDA. In addition, postmortem immunohistochemistry against TH revealed less severe neurodegeneration of DA neurons compared to vehicle-injected control animals. Furthermore, when mice were pretreated with ANA-12, a selective antagonist for tropomyosin receptor kinase B (TrkB), the behavioral improvement and neuroprotective effect were diminished. These results suggest that increased neuronal activity of DA neurons provides neuroprotection against 6-OHDA injury and alleviates its symptoms through the brain-derived neurotrophic factor-TrkB pathway.

## Introduction

Parkinson’s disease (PD) is the second-most common neurodegenerative disorder. Degeneration of dopaminergic (DA) neurons in substantia nigra pars compacta (SNC) worsens motor symptoms over time (Lees et al., 2009). Tremendous efforts have been directed towards suppressing or slowing down this continuous degeneration. Studies showed that the signaling pathway involving brain-derived neurotrophic factor (BDNF) and its downstream receptor tyrosine kinase (TrkB) have pro-survival effects (Nagahara and Tuszynski, 2011). In SNC of patients with PD, a reduction of BDNF expression in surviving DA neurons was observed (Howells et al., 2000). Therefore, enhancing the BDNF-TrkB signaling pathway can be a potential therapy for PD (Nagahara and Tuszynski, 2011). As BDNF is unable to cross the blood brain barrier (BBB), invasive surgery is necessary to deliver BDNF in the gene or protein form directly to the brain (Allen et al., 2013). Rodent and primate studies of PD demonstrated promising results (Tsukahara et al., 1995; Hernandez-Chan et al., 2015). However, as a surgery is generally irreversible, it is difficult to manipulate the level of BDNF after the surgery. Systemic injection of TrkB agonists also showed neuroprotective results in PD models(Jang et al., 2010; Nie et al., 2015). However, the lack of specificity may cause unintended wide-spread activation of TrkB, potentially resulting in serious side effects such as seizures (Binder et al., 2001).

BDNF does not necessarily have to be delivered exogenously. An *in situ* hybridization study has shown that endogenous BDNF is widely distributed in the human brain, including DA neurons in SNC (Conner et al., 1997; Howells et al., 2000). BDNF is released from neurons via the constitutive secretory pathway as well as the activity-dependent secretory pathway (Park and Poo, 2013). Electrical stimulation that increases neuronal activity can lead to BDNF release *in vitro* (Al-Majed et al., 2000). *In vivo* studies in rodents have further shown that significant contribution of BDNF in the neuroprotective effects mediated by physical exercise (Spieles-Engemann et al., 2010) as well as by deep brain stimulation in the subthalamic nucleus (DBS-STN) (Spieles-Engemann et al., 2010; Spieles-Engemann et al., 2011). A TrkB antagonist diminished the neuroprotective effects by physical exercise or DBS-STN (Real et al., 2013; Fischer et al., 2017a), indicating involvement of the BDNF-TrkB pathway. Epidemiologically, people who exercise often have lower risk of PD (Xu et al., 2010). Therefore, enhancement of endogenous BDNF release may be an effective neuroprotective therapy for PD. To this end, conventional optogenetics has been utilized to activate the BDNF-TrkB pathway *in vitro* (Park et al., 2015) and *in vivo* (Carreno et al., 2016). Such an approach has been deployed to promote regeneration after stroke (Cheng et al., 2014).

In this study, in order to provide prolonged neuromodulation to DA neurons more conveniently and less invasively, we employed bioluminescence-optogenetics using step-function luminopsin (SFL) that our group had developed previously (Berglund et al., 2020). SFL consists of *Chlamydomonas* channelrhodopsin 2 (CrChR2) with double point mutations (C128S/D156A)(Yizhar et al., 2011b) and *Gaussia* luciferase (GLuc) with double point mutations (M60L/M127L)(Welsh et al., 2009). In the presence of GLuc substrate, coelenterazine (CTZ), slow-deactivating step-function ChR2 is activated by improved bioluminescence from GLuc, led to sustained depolarization in neurons. DA neurons was targeted through stereotactic injection of recombinant adeno-associated viral (AAV) vectors carrying the SFL gene into SNC.

As a mouse model of PD, unilateral injection of the DA-depleting neurotoxin, 6-hydroxydopamine (6-OHDA), provides several benefits including low mortality, using the unlesioned side as an internal control, and a readily quantifiable behavior (ipsiversive rotations)(Blesa et al., 2012). One drawback of this model, however, is the acute nature of the effect, which diminishes over time and allows mice to recover from the behavioral deficit (Iancu et al., 2005; Thiele et al., 2012; Bagga et al., 2015; Boix et al., 2015). Therefore, we first optimized the dosage and injection site of 6-OHDA for a long-term study up to one month.

We further demonstrated that chronic activation of DA cells through SFL and daily injection of CTZ attenuated behavioral deficits and neuronal loss. Furthermore, TrkB blockade abolished this effect, suggesting involvement of the BDNF-TrkB signaling in this activity-dependent neuroprotection.

## Materials and methods

### Plasmids

DNA plasmids were made using conventional molecular biology methods and restriction enzymes (RE; New England BioLabs, Ipswich, MA). An adeno-associated virus serotype 2 (AAV2)-based transfer vector with the human synapsin I promoter (hSyn) and step-function luminopsin (SFL) was described previously(Berglund et al., 2020). To enable Cre recombinase-dependent expression, the NheI – BsrGI fragment containing SFL in the pcDNA3.1/CAG vector was ligated into the corresponding RE sites of an AAV2 vector with the elongation factor 1α promoter and double-floxed inverted open reading frame (EF1a-DIO; a gift from Karl Deisseroth; Addgene plasmid #: 55631; RRID: Addgene_55631) in the untranslatable reverse orientation. To reduce the size of cargo, the polyadenylation terminator was replaced with a shorter one from the bovine growth hormone gene by ligating the EcoRI – RsrII fragment from another EF1a-DIO vector (a gift from Ute Hochgeschwender; Addgene plasmid #: 114105; RRID: Addgene_114105). The correct modifications were confirmed by RE analyses and DNA sequencing. Endotoxin-free, transfection-grade plasmids were produced using midi-prep kits (NucleoBond Xtra Midi EF; Macherey-Nagel, Düren, Germany).

### Viral vectors

Recombinant AAV vectors pseudotyped with AAV9 were produced in Emory University Virus Core. Viral particles were resuspended in sterile PBS, aliquoted, and kept at -80°C until use. Titers were determined by quantitative PCR for the inverted terminal repeats. They were 1.0 × 10^13^ viral genomes (vg)/ml for hSyn-SFL and 1.6 × 10^13^ vg/ml for EF1a-DIO-SFL.

### Animals

All procedures involving mice were conducted in accordance to protocols approved by the Emory University Institutional Animal Care and Use Committee. Wildtype male mice aged between 8 and 11 weeks were purchased (strain: C57BL/6J; stock #: 00664; Jackson Laboratory, Bar Harbor, ME). Hemizygous transgenic male mice were bred inhouse by crossing BAC-transgenic mice expressing bacteriophage Cre recombinase under control of the mouse tyrosine hydroxylase promoter (TH-Cre; MMRRC stock #: 017262-UCD; RRID: MMRRC_017262-UCD) with wildtype mice. They were genotyped according to the repository’s PCR protocol using biopsy samples and the strain-specific primers. The transgenic mouse colony had been maintained on a C57BL/6J background for at least 6 generations. Animals were housed in the Emory University animal facility with *ad libitum* standard rodent chow and water with a 12-hour light/12-hour dark cycle.

### Surgeries

Mice were anesthetized with 1.5-3% isoflurane in pure oxygen and placed into a stereotactic frame (David Kopf Instruments). One microliter of AAV (∼10^10^ viral particles) was stereotactically injected into substantia nigra pars compacta (SNC) through a burred hole in the skull using a pulled glass pipette and a microinjector (Nanoject; Drummond Scientific) at a rate of 275 nl/minute. The pipette was removed 5 minutes after injection and the scalp was glued (VetBond, 3M). The coordinate for injection was 3.1 mm posterior and 1.6 mm lateral from the bregma and 4 mm below the dura. In some animals, a multielectrode array (MEA; a 4 by 4 square array with tungsten electrodes of 6 mm long and 35 µm in diameter with 150 µm between-wire spacing; Innovative Neurophysiology, Durham, NC, USA) and an optical fiber ferrule (10 mm with 200 µm core diameter and 0.39 numerical aperture (NA); R-FOCL200C-39NA, RWD) was implanted stereotactically after virus injection. The fiber ferrule was attached to the MEA with VetBond (Tung et al., 2015). The coordinate for MEA implantation was 3.4 mm posterior and 1.25 mm lateral from the bregma and 4.7 mm below the dura. The MEA was affixed to the skull with dental cement (Ortho-Jet, Lang Dental). 20 days after the virus injection, 6-hydroxydopamine (6-OHDA; cat #: 162957; Sigma-Aldrich) was injected similarly into the right striatum ipsilateral to the virus injection. The neurotoxin was freshly dissolved immediately before surgery at a concentration of 2.67 mg/ml in 0.9% (w/v) NaCl sterile saline supplemented with 0.1% (w/v) ascorbic acid (cat #: A4403; Sigma-Aldrich) and kept from sunlight on ice until injection. The mouse was monitored daily after surgery. When a significant loss of body weight was observed, mice were handfed. All the mice maintained body weight above the removal criterion of 20% loss.

### Intraperitoneal injections

SFL substrate, native coelenterazine (CTZ; cat # 303-INJ; Nanolight Technology) was freshly reconstituted in sterile proprietary solvent (NanoFuel; 500 µg/100 µL). CTZ was injected 4 days before the 6-OHDA surgery. The injection is daily except the day that need to perform rotation test. The control group was injected only with the solvent.

In order to block the BDNF-TrkB pathway, ANA-12 (cat #: 506304; Millipore Sigma) was used. The selective TrkB antagonist was dissolved in a solution consisting of 66% Dulbecco’s phosphate-buffered saline (PBS), 16.5% ethanol, 16.5% CremophorEL, and 1% dimethyl sulfoxide (in v/v). Intraperitoneal injection of ANA-12 (0.5 mg/kg) was performed 4 hours before each CTZ/vehicle injection, when the effect of ANA-12 is expected to be maximized (Cazorla et al., 2011).

### Behavioral test

A mouse was put into a glass cylinder (diameter: 11 cm). A baseline video recording started 5 minutes after placement and lasted for 30 minutes. The mouse then received intraperitoneal injection of amphetamine (2.5 µg/kg) dissolved in sterile saline. Another video recording under amphetamine started 5 minutes after injection and lasted for 45 minutes. Rotation assays were performed one day before 6-OHDA injection and once a week afterwards for 2 weeks for wildtype mice injected with hSyn-SFL and for 4 weeks for TH-Cre mice injected with EF1a-DIO-SFL. Full ipsiversive and contraversive rotations were counted and the numbers of net ipsiversive rotations (the number of ipsiversive rotations subtracted by that of contraversive ones) were used for the analysis. The scorers were blinded to the experimental conditions and groups.

### Electrophysiology, optogenetics, and fiber photometry

Extracellular multi-unit recordings were conducted at 100 kHz using a 32-channel recording system (Tucker-Davis Technologies) under isoflurane anesthesia (2 – 3 % in oxygen). Single unites were isolated offline using a free software (KiloSort). Photostimulation was delivered through the implanted fiber ferrule connected to two fiber-coupled LEDs (465 nm and 590 nm; Plexon). Irradiance was measured at the tip of the fiber as 100 and 95 mW/mm^2^, respectively. The two LEDs were fed to the same fiber ferrule through a light spectrum mixer (LSM_1×3_470/530/590_FC, Doric). For bioluminescence fiber photometry, the same fiber ferrule was connected to a high-sensitivity amplified photodiode (PDF10A/M; Thorlabs) through a patch cable (core diameter: 200 µm; NA: 0.39; Thorlabs). The signal was sampled at the same time as multi units in an analog channel of the electrophysiology system. The recording consisted of 15-s baseline, 1-s 465 nm stimulation, followed by 20-s no-light period. The activation of SFL was reversed by illuminating 590 nm light for 3 s.

### Histology

After completion of the final rotation assay, the animals were sacrificed by intraperitoneal injection of Euthasol followed by cardiac perfusion with 4% (w/v) paraformaldehyde in saline. The brains were dissected and post-fixed in the same solution at 4°C overnight. Then, the brains were cryoprotected by 30% (w/v) sucrose in PBS at 4°C overnight. Coronal sections were cut 20 µm thick for immunofluorescence and 40 µm thick for immunohistochemistry from frozen brains using a cryostat. For immunofluorescence, free-floating sections were permeabilized and blocked with 0.1% (v/v) Triton X-100 and 4% (v/v) normal goat serum in PBS at room temperature for 30 minutes and then incubated with a rabbit anti-TH antibody (Pel-Freeze, cat # P40101; 1:1,000 dilution) at 4°C overnight. Then, the sections were incubated at room temperature for 2 hours with a secondary antibody: goat anti-rabbit IgG antibody conjugated with Alexa Fluor 594 (Invitrogen, cat # A-11012; 1:1,000 dilution). For immunohistochemistry, sections were permeabilized with 3% (v/v) hydrogen peroxide and 0.1% (v/v) Triton X-100 in PBS for 20 minutes. The sections were rinsed with PBS for 10 minutes 3 times. The sections were then blocked with 4% (v/v) normal donkey serum in PBS for 30 minutes. Then, they were incubated with the same primary antibody in 2% (v/v) normal donkey serum in PBS at 4°C overnight. The sections were washed with PBS for 10 minutes 3 times and incubated with a biotinylated secondary antibody followed with avidin– HRP solution (Vectastain ABC HRP Kit; cat #: PK-4000; Vector Laboratories, Burlingame, CA) at 4°C overnight. After 3 washes for 30 minutes each with PBS, the sections were then developed in 3-39-diaminobenzidine tetrachloride for 2–3 minutes. After several rinses with PBS, sections were mounted onto glass slides, dehydrated, and coverslipped.

### Microscopy

To quantify the number of TH-immunopositive DA neurons, cell body counting was performed using an upright microscope (Nikon Eclipse E800) equipped with a CCD camera (JVC). TH-positive neurons were counted in both ipsilateral and contralateral SN and the adjacent ventral tegmental area (VTA) (Carman et al. 1991). The stereotaxic region to be examined in each section was determined by the shape and distribution of TH-positive neurons at lower magnification (40x) first with reference to the standard mouse brain atlas (Paxinos and Franklin, 2001). Cells with intact nuclei, clear cytoplasm, and neuronal processes were included in the counts and cells with deformed cell bodies and no obvious processes were excluded from the counts (Iravani et al., 2002). Sections were counted at each stereotaxic level and these were pooled to give a mean cell count for ipsilateral and contralateral hemispheres in each group. The observers were blinded to the experimental conditions.

### Statistical analysis

Prism v8.1.2 software (GraphPad, San Diego, California) was used for statistical analyses. Two-way ANOVA followed by Tukey’s multiple comparisons tests was used to compare the spontaneous and amphetamine rotations. For neuronal count, an unpaired two-tailed *t*-test was used to compare TH staining. All results are presented as mean ± SEM.

## Results

### Characterization of 6-OHDA lesion models

First, we evaluated different doses and locations of unilateral 6-OHDA injections through behavioral and pathological changes. We tested three dosages (1.5, 2.67, or 3 µg) in the striatum (STR) and one dose in the medial forebrain bundle (MFB; 3 µg). We observed increases in ipsiversive rotations for up to 1 month after injections of 2.67 µg or 3 µg 6-OHDA in the STR (**Fig. 1A**). Two-way ANOVA with repeated measures revealed significances in the main effect of dosage (*F*(2, 13)=4.431; \**p*=0.034; *n*=5, 6, and 5 mice for 6-OHDA 1.5, 2.67, and 3 µg, respectively) as well as that of time points (*F*(4, 52)=14.83, \*\*\**p*<0.0001) and their interactions (*F*(8, 52)=4.175, \*\*\**p*=0.0007). With the dosage of 2.67 µg, the numbers of ipsiversive rotations at all the time points tested after 6-OHDA injection were significantly larger than the pre-injection level (\*\*\**p*<0.0001 for all the timepoints; *q*<0.0001 for all the time points; Tukey’s multiple comparisons tests) while with the dosage of 3 µg, significant increases were only observed at 14, 21, and 28 days post injections (\**p*=0.007, 0.0197, and 0.031; *q*=0.024, 0.036, and 0.039, respectively). We observed a similar but smaller effect of the MFB injections on ipsiversive rotations. A significant difference was not observed in animals with the dosage of 1.5 µg 6-OHDA (*p*>0.2; *q*>0.99 for all the time points).

**Figure 1.**
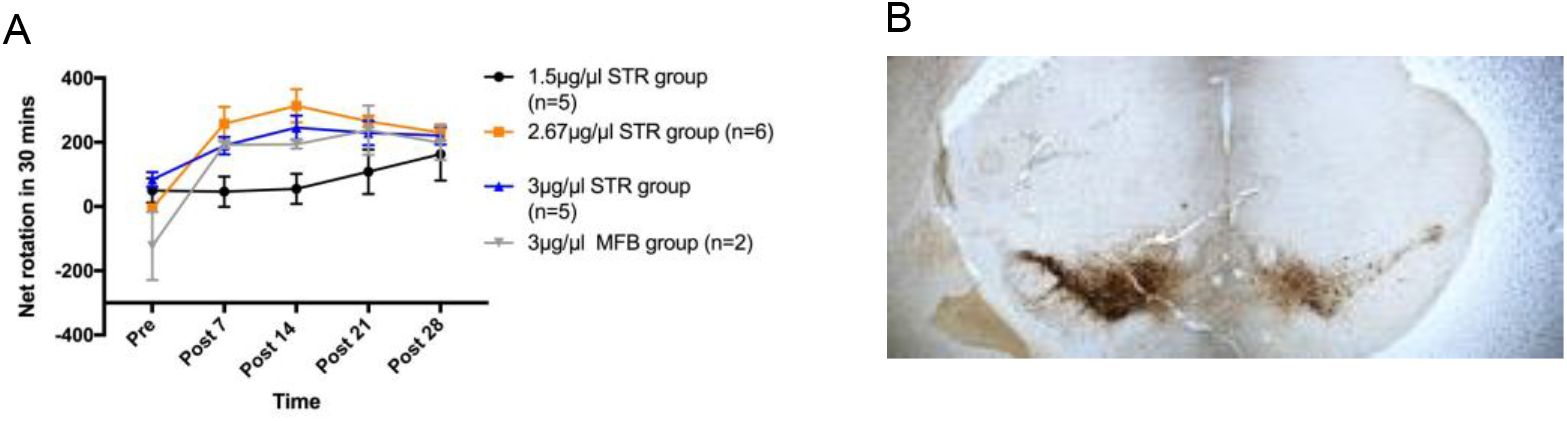
The effect of 6-OHDA with different dosages and injection sites. A) We observed significant amphetamine induced rotation effect in 2.67 ug/ul STR group on every timepoint. We also observed significant but less robust rotation increased in 3ug/ul STR group, and 3ug/ul MFB group. B) Representative image of TH-DAB immunohistochemistry against TH. 6-OHDA was injected into the right striatum at 2.67ug/ul.

We further characterized pathological changes in the SNC after 6-OHDA injection (2.67 µg) in the STR by conducting immunostaining against TH, a marker for DA neurons. Reduction of TH immunoreactivity was observed in the 6-OHDA-injected side (**Fig. 1B**), consistent with loss of DA neurons in the SNC. Based on the above results, we used unilateral injection of 2.67 µg of 6-OHDA in the STR for the rest of this study.

### Pan-neuronal SFL expression and CTZ reduced behavioral abnormality but not neuronal loss

We first injected an AAV vector with the pan-neuronal human synapsin I (hSyn) promoter into substantia nigra of wildtype mice to examine the effect of activation of neurons through non-specific expression of SFL and systemic CTZ administration in the 6-OHDA model. Two weeks after virus injection, we started pretreatment with either CTZ (10 mg/kg) or vehicle through intraperitoneal injections 4 days prior to the 6-OHDA injection. The treatment with CTZ or vehicle continued 6 days a week for two weeks until the end of experiments, when the animals were sacrificed for histological analysis (**Fig. 2A**). hSynI-driven expression of SFL (**Fig. 2B**) was observed through the enhanced yellow fluorescent protein (EYFP) tag in neurons around the injection site, including SNC and substantia nigra pars reticulata (SNR) (**Fig. 2C**). We quantified the number of ipsiversive rotations, the behavioral abnormality caused by 6-OHDA, at two different time points, 1 day before and 14 days after 6-OHDA injection, in two different settings, spontaneous rotations and rotations observed under the influence of amphetamine. Two-way ANOVA with repeated measures revealed significances in the main effect of CTZ vs. vehicle in spontaneous rotation (*F*(1, 13) = 4.732; \**p*=0.0487; *n* = 8 and 7 mice for vehicle and CTZ, respectively). Spontaneous rotations in the CTZ treatment group were significantly lower than the vehicle treatment group at day 14 (\**p* = 0.0136; *q*=xx**Fig. 2D)**. However, there was no significant difference in the main effect of CTZ in rotations observed under amphetamine between the two groups (F (1, 13) = 2.850; *p* = 0.1152; **Fig. 2E**). We did not observe a significant difference in neuronal loss in SNC between the CTZ- and vehicle-treated mice (*p* = 0.142; **Fig. 2F**). As a control for the CTZ treatment, another set of animals received injection of a control vector carrying the GFP gene only that did not react with CTZ. Unlike the mice with SFL, we did not observe a significant difference between the CTZ and vehicle groups in the rotational assays (for spontaneous rotation *F*(1, 11) = 0.9460, *p* = 0.3516; for amphetamine rotation *F*(1, 11) = 1.369, *p* = 0.2667; **Fig. 2G & H**).

**Figure 2.**
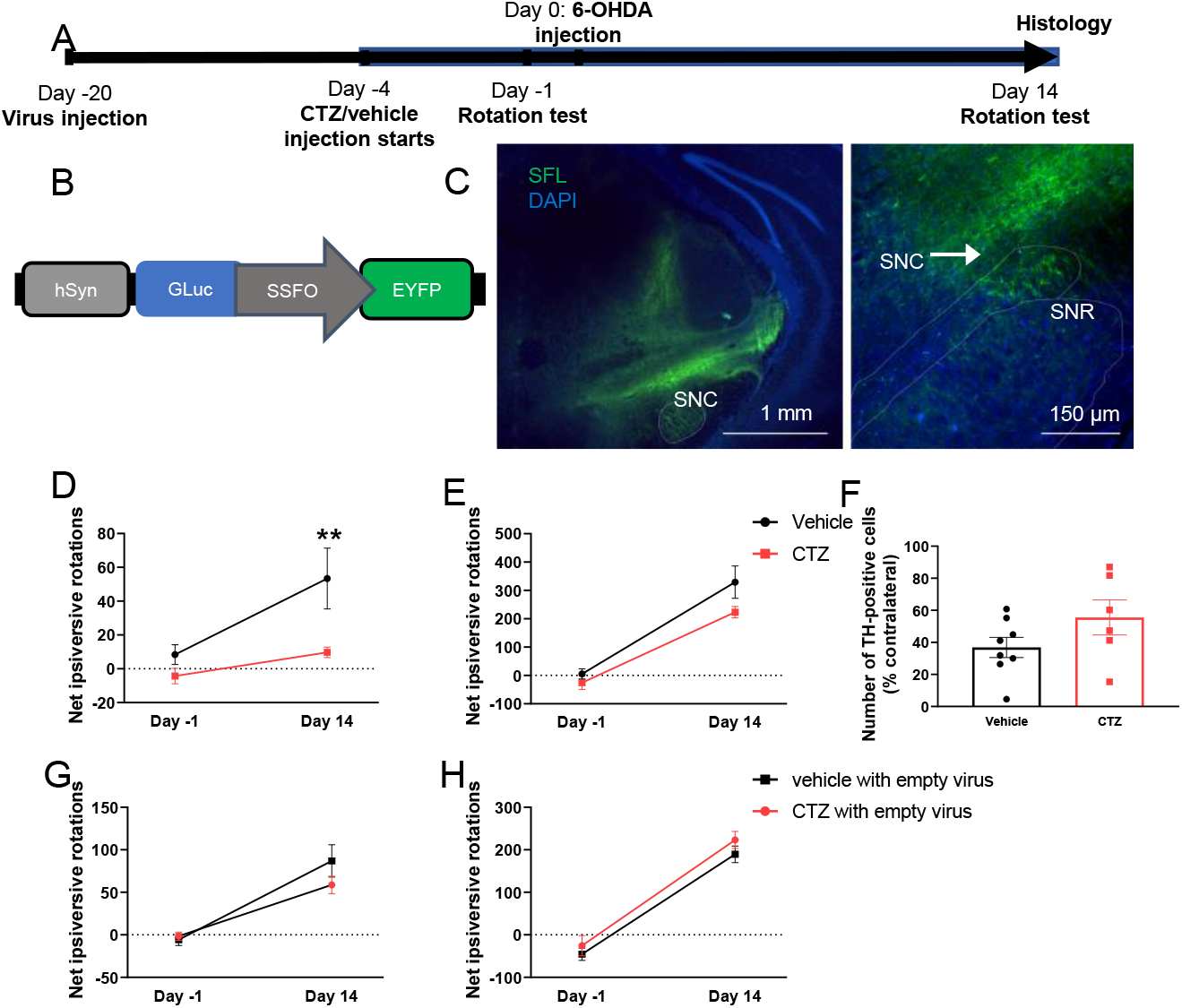
Expression is nonselective in hysn-SFL virus in wild type mice. SFL reduced only ipsiversive spontaneous rotation and does not prevent neuronal loss in wild type mice. A) Experimental timeline of wild type mice group. B) Schematic of the hSyn-SFL construct. C) Virus expression spread everywhere around the SNC region. D) SFL was able to reduce ipsiversive spontaneous rotation in CTZ compared vehicle at day 14. E) Amphetamine rotation between CTZ and vehicle were not significantly changed. F) No significant changes in comparing SNC neurons between CTZ and vehicle. G-H) No significant changes in spontaneous and amphetamine rotation for GFP only virus.

### Cre-dependent viral vector for conditional expression of SFL

In order to achieve cell type specificity, we constructed an AAV vector (DIO-SFL) for Cre-dependent expression of SFL. Cre-dependent expression of SFL was confirmed using a heterologous expression system (human embryonic kidney 293 cells) and lipofection. The mammalian cell line was transfected with the DIO-SFL plasmid together with a Cre plasmid and SFL expression was examined as fluorescence from the EYFP tag (**Supplementary figure 1**). SFL expression was observed only in cells transfected with both plasmids, but not in cells with the Cre plasmid or the DIO-SFL plasmid alone. As a positive control, cells were transfected with a plasmid with the SFL gene in the normal orientation, resulting in similar expression of SFL primarily on the membrane. These results confirmed Cre-dependent expression of SFL using the constructed plasmid.

An AAV vector pseudotyped with AAV9 was produced with this transfer plasmid and was injected in SNC in a transgenic mouse line with expression of Cre under control of the TH promoter (TH-Cre). To optimize expression of Cre-dependent expression of SFL, we injected two dosages (1.6×10^9^ or 1.6×10^8^ vg). As a negative control, we injected the higher dosage into a TH-Cre-negative littermate as well. Two weeks after injection, we sacrificed the animals and prepared conventional histological sections for observation with fluorescence microscopy. We observed a specific pattern and a suitable level of SFL expression in the SNC with the higher dosage (**Supplementary figure. 2a**), but not with the lower dosage (**Supplementary figure 2b**). No visible expression was seen in the negative control mouse (**Supplementary figure 2c**).

We further characterized neuromodulation of the SNC by SFL through extracellular recordings under isoflurane anesthesia (**Fig. 3**). SFL enables bi-modal neuromodulation through conventional physical light and CTZ-activated biological light (Berglund et al., 2020). We typically encountered two different types of cells: presumable DA cells with wide action potential waveform at low frequency and presumably GABAergic cells with narrow action potential waveform at high frequency (**Figs. 3A & B, respectively**). Both of the cell types responded to photostimulation of 465 nm light of LED and the response was reversed by longer wavelength of light (590 nm), but to the opposite direction: Whereas we typically observed an increase of action potential firing in a putative dopaminergic cells during photostimulation, we did a decrease in a putative GABAergic cell (**Figs. 3C & D, respectively**). Overall, about one third of cells recorded responded to photostimulation with a majority cells of slow spiking (<10 Hz; **Fig. 3E**). A majority of cells with less than 10-Hz spiking showed increases of firing whereas cells with more than 10-Hz spiking showed decreases (**Fig. 3F**). Further, we observed an increase of firing after intraperitoneal injection of CTZ (20 mg/kg body weight), concomitantly with an increase of bioluminescence collected through the same fiber ferrule used for photostimulation with external light (**Supplementary figure 3**). These results are consistent with proper targeting of DA neurons with excitatory SFL and showed its ability to modulate neuronal firing through external and internal light.

**Figure 3.**
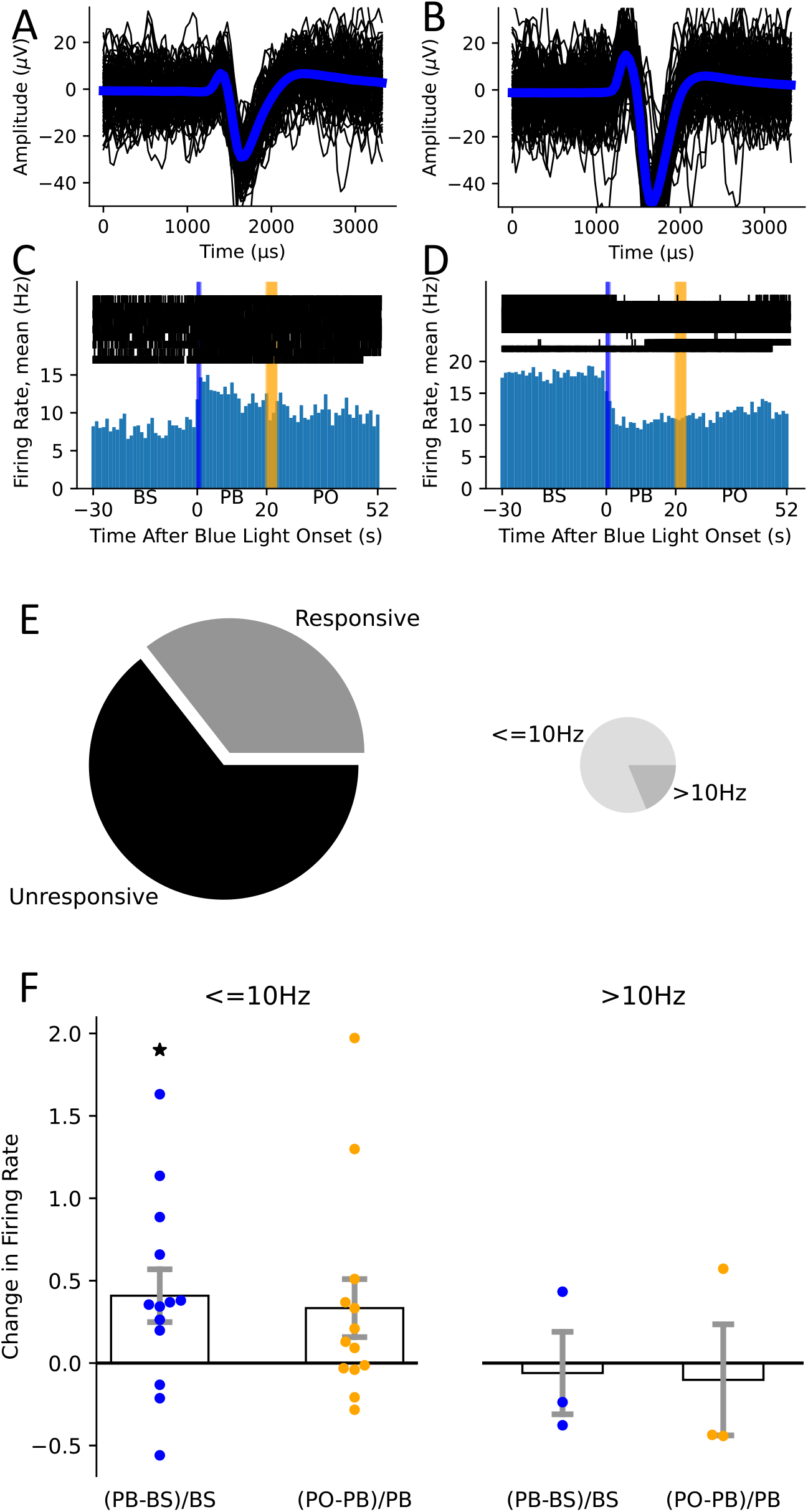
Effects of SFL activation on single-unit electrophysiology in the SNC. Each trial consists of a period before stimulation (BS), brief stimulatory (blue) light followed by a post-blue (PB) period, then a pulse of inactivating (orange) light and a post-orange (PO) period. Units (example waveforms shown in A, B) exhibited a mixture of non-response, excitation (C), and inhibition (D) in response to blue light. The population of responsive units could be further divided into units with slow (<=10Hz) and fast (>10Hz) baseline activity (E). Only the slow units showed a significant difference in their pre and post blue light firing rate (\**p*=.0254, paird *t*-test).

### Prolonged activation of DA neurons via SFL and CTZ reduced ipsiversive rotations and neuronal loss

To test neuroprotection through chronic and prolonged activation of DA cells in the PD model, we injected the DIO-SFL vector into the SN of TH-Cre mice (**Fig. 4A**). Compared to the hSyn vector, expression of SFL was more limited in the SNC and appeared to be in DA cells only (**Figs. 4B & C)**. TH-Cre mice with DIO-SFL were subjected to daily treatments of CTZ or vehicle 4 days prior to the 6-OHDA surgery and, after the surgery, 5 days a week for four weeks until they were sacrificed (**Fig. 4A**). Daily CTZ administration led to a significant reduction of ipsiversive rotations (**Figs. 4E & F**): We observed significant differences in the two groups in both spontaneous (*F*(1, 12) = 16.78; \*\**p* = 0.0015; *n* = 8 and 6 for the control and CTZ group, respectively) and amphetamine-induced rotations (*F*(1, 12) = 17.08, \*\**p* = 0.0014; *n* = 8 and 6 for the control and CTZ group, respectively). Multiple comparisons between the two groups at different time points revealed significant differences in the amphetamine condition 7 days after 6-OHDA injection (\*\**p* = 0.0035). At 14 and 21 days after 6-OHDA injection, there was significant reduction in both spontaneous (\**p* = 0.0002 and 0.0239, respectively) and amphetamine-induced rotations (\*\*\**p* = 0.0001 and 0.0003, respectively).

**Figure 4.**
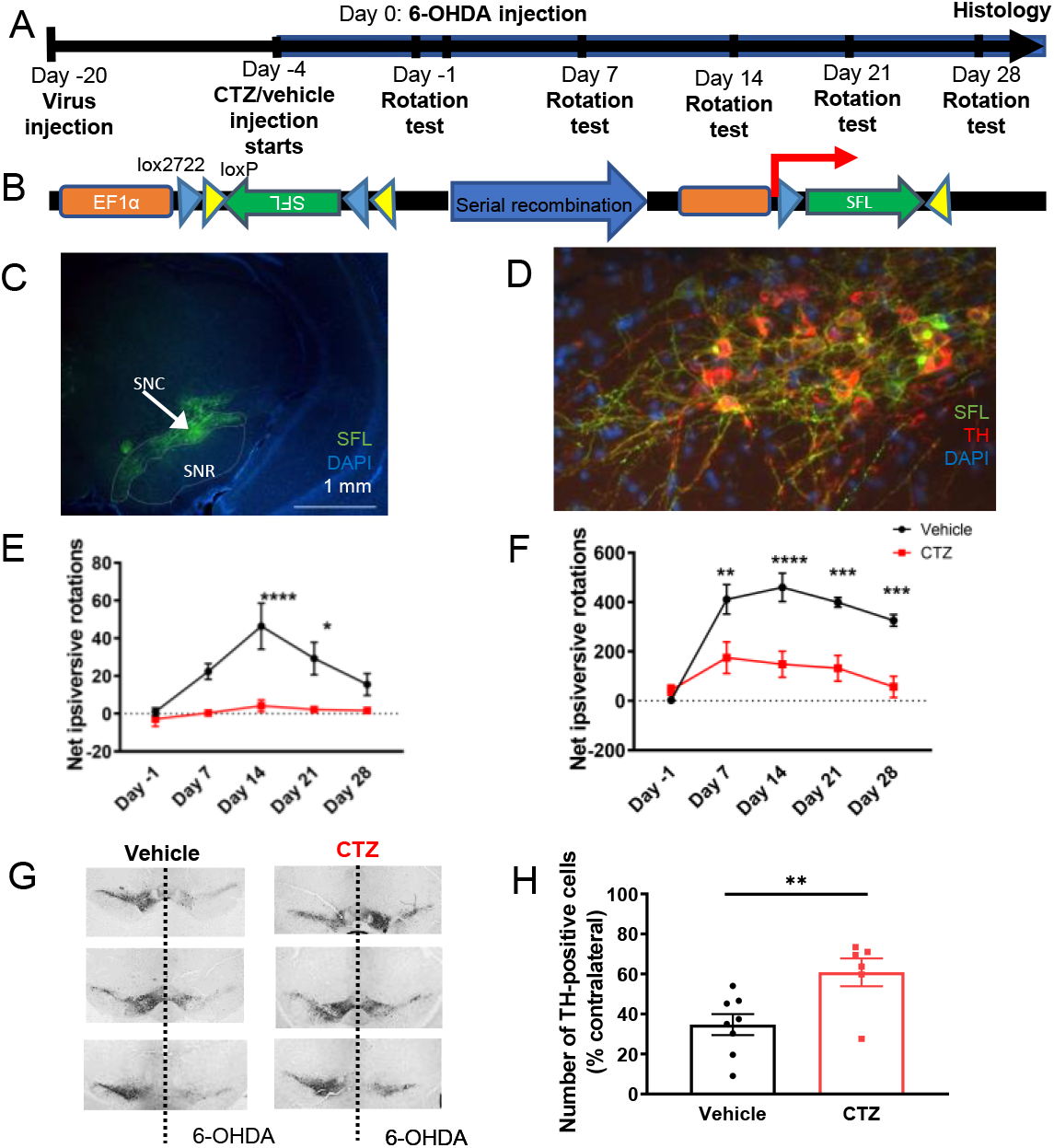
Expression is selective in DIO-EF1α-SFL virus in TH+ mice. There are significant improvement of survival of TH-expressing neurons with the CTZ treatment. SFL reduced ipsiversive spontaneous and amphetamine rotations and prevent neuronal loss in TH-Cre mice A) Experimental timeline of TH+ type mice group. B) Schematic of the hSyn-SFL construct. C) Virus expression mainly target in TH+ neurons in SNC. D) SFL was able to reduce ipsiversive spontaneous rotation in CTZ compared vehicle from day 7 to day 28. E) SFL was able to reduce ipsiversive amphetamine rotation in CTZ compared vehicle from day 7 to day 28. G) Immunoperoxidase staining for TH+ on representative coronal midbrain sections showing less SNC TH + neurons on vehicle compared to CTZ. H) The neurodegenerative effect was lessened by the CTZ treatment. \**p* < 0.05; two-tailed *t*-test; *n* = 8 and 6 mice for vehicle and CTZ, respectively.

To further elucidate the neuroprotective effects by SFL and CTZ, we examined pathological changes in SNC (**Fig. 4G**). Neurodegeneration of TH-immunopositive neurons in SNC was confirmed by loss of TH staining in the 6-OHDA injected side. Consistent with the behavioral effect, TH staining in the 6-OHDA injected sides were relatively stronger in the CTZ-treated animals than in the vehicle-treated animals (control). On average, TH-positive, surviving neurons in SNC was significantly higher in the CTZ-treated animals than the vehicle group (*t*(12) = 3.069, \**p* = 0.0097; *n* = 8 and 6 mice for control and CTZ, respectively; **Fig. 4H**). This immunohistochemical result is consistent with the effect of CTZ observed in the behavioral paradigm, suggesting that survival of TH-expressing neurons improved by the CTZ treatment underlies the better behavioral outcome.

### Neuroprotective effect of SFL and CTZ on DA cells was partially mediated by BDNF/TrkB pathway

Previous research showed increased neuronal activity and activation of the BDNF/TrkB pathway can lead to neuroprotective effects. In order to investigate whether the neuroprotective effects through SFL and CTZ were mediated through BDNF, we used ANA-12, a selective antagonist of TrkB receptor. ANA-12 (500 µg/kg) was administered 4 hours prior to daily treatments with CTZ or vehicle. When pretreated with ANA-12, ipsiversive rotations in the CTZ group appeared similar to those in the vehicle control group (**Figs. 5C & D**) and, in fact, there was no statistical difference between the two groups (for spontaneous rotation, *F*(1, 11) = 0.03229, *p* = 0.8606; for amphetamine rotation, *F*(1, 12) = 0.48, *p* = 0.5016). When the CTZ alone group was compared to the CTZ plus ANA-12 group, we observed a significant reduction in spontaneous ipsiversive rotations (\**p*= 0.0398; the main effect of treatment; two-way ANOVA with repeated measures; *F*(1,14) = 5.139 ;*n* = 6 and 8 mice for CTZ and CTZ+ANA-12, respectively**; Fig. 5E)**. In the amphetamine-induced rotations, the results between animals treated with CTZ plus ANA-12 and CTZ alone were similar and, in fact, the main effect of treatments was not significant (*F*(1,14) = 0.07505; *p* > 0.7; **Fig. 5F**). Taken together, these results indicate that ANA-12 abolished the neuroprotective effect of CTZ, at least on spontaneous ipsiversive rotations.

**Figure 5.**
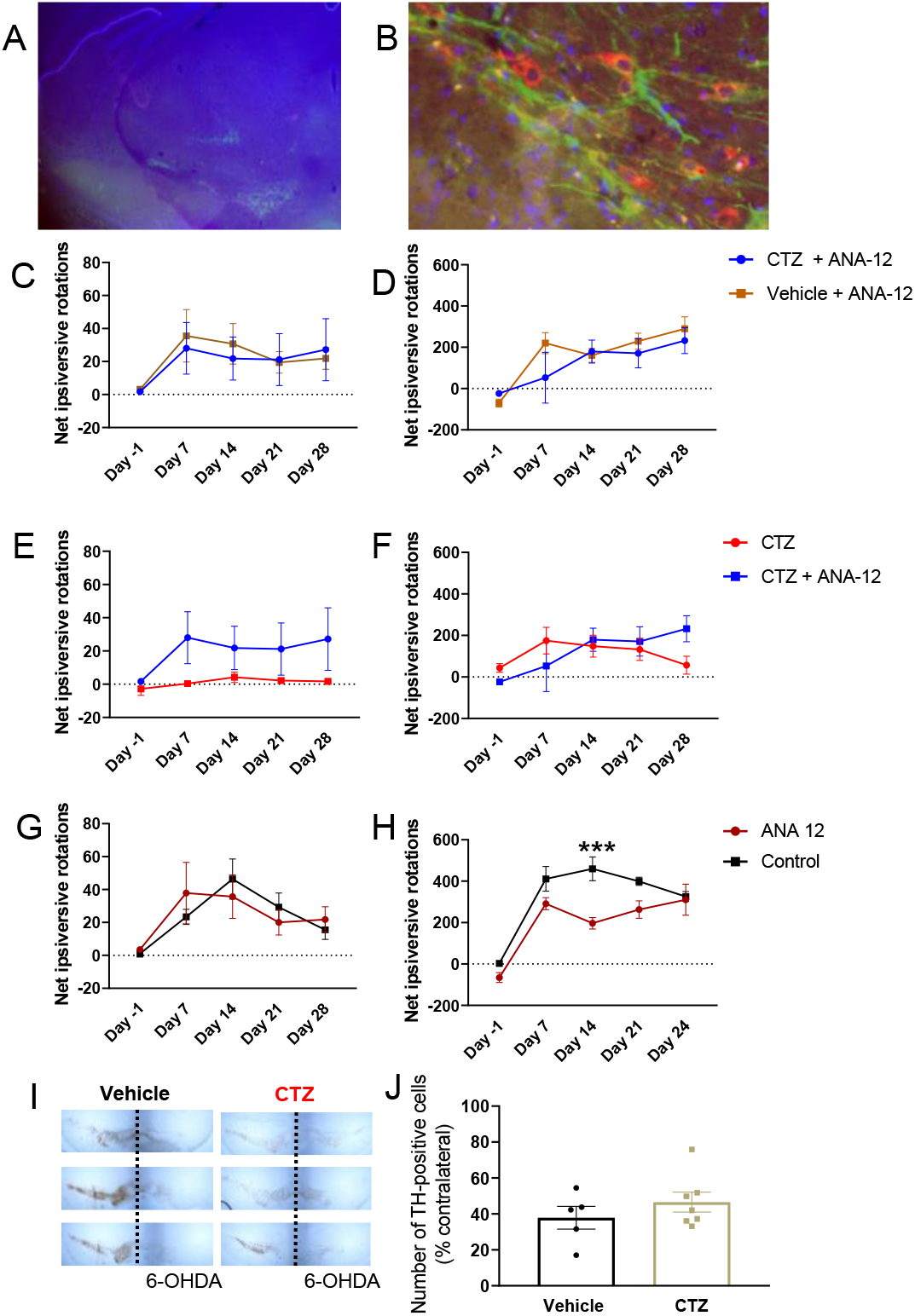
ANA-12 diminished spontaneous rotation recovery and increased of neuronal survival induced by CTZ. A-B) Virus expression mainly target in TH+ neurons in SNC. C-D) With ANA-12, there were no significant change of spontaneous and amphetamine rotation between CTZ and vehicle group. E)A significant increase of spontaneous rotation in CTZ+ANA-12 compared to CTZ+vehicle. F) No significant change of amphetamine rotation in CTZ+ANA-12 group compared to CTZ+vehicle group. G) No significant change of spontaneous rotation in ANA-12 compared to control. H) A significant reduction in amphetamine rotation in ANA-12 group compared to control group. I) Immunoperoxidase staining for TH+ on representative coronal midbrain. J) No significant difference in neuronal loss in ANA-12 - vehicle and ANA-12-CTZ group.

We also examined the effect of ANA-12 by itself without the CTZ treatment. We did not observe any exacerbation of the 6-OHDA injury by daily treatments of the TrkB blocker: There was no significant difference in spontaneous ipsiversive rotations (*F*(1,16) = 0.6312; *p* > 0.5; **Fig. 5G**) and, under the amphetamine condition, we rather observed a significant reduction of ipsiversive rotations (*F*(1,16) = 16.05; \*\*\**p* = 0.001; **Fig. 5H**). These results indicate that the BDNF/TrkB pathway was not active to exert an endogenous neuroprotective effect. We further examined pathological changes in the SNC through immunohistochemical staining (**Fig. 5I**). On average, there was no significant difference in TH-positive neurons in SNC between the CTZ and vehicle group when the animals were pretreated with ANA-12 (*p* = 0.33; **Fig. 5J**). As ANA-12 blocks the TrkB pathway, our results suggest that the neuroprotective effect we observed with SFL and CTZ was due to activity-dependent release of BDNF and subsequent activation of the TrkB pathway.

## Discussion

Our study shows that selective, chronic activation of DA neurons can increase neuronal protection and reduce behavior deficits. In contrast, this protective effect is relatively modest when virus is not cell-type specific. There was no behavioral recovery when GFP only virus was used. This shows the recovery effect is due to increase neuronal activity caused by luminopsin. In contrast, Th-Cre mice model of PD with CTZ injection showed reduced ipsiversive rotations and had higher neuronal count. This highlights the importance of transducing the right cell type. One possible explanation for this disparity is that not all DA neurons were being transduced in wild type mice which limited the protective effect. Another possibility is that virus could be transducing neurons that inhibit DA neurons. It has been shown there is an inhibitory interaction between GABAergic neurons in SNR and DA neurons (Tepper et al., 1995).

Our results also demonstrate the important role of BDNF/TrkB signaling. Spontaneous rotations and neuroprotection were abolished when ANA-12 was given before CTZ injection. Interestingly, amphetamine rotation remained the same. When comparing ANA-12 plus vehicle group with just vehicle group, there was a significant reduction of ipsiversive rotations only in amphetamine-induced rotations, but not in spontaneous rotation, indicating a possible interaction between amphetamine and the BDNF-TrkB pathway (Shen et al., 2006). Our finding suggests that neuroprotection was mediated by BDNF-TrkB signaling pathway. Our study demonstrates that the neuroprotective effect can be obtained only when activating DA neurons, and the effect diminished when SFL expression was non-selective through the pan-neuronal hSyn promoter.

Compared to previous treatments with BDNF gene/protein or general neurotrophic factors, our study provides several advantages. Our treatment is capable of increasing the BDNF level in the brain in a controllable manner compared to gene therapies with neurotrophic factors, which are permanent and irreversible. We can control the secretion of BDNF based on the dosage and frequency of CTZ administration, which is difficult for the surgery and gene therapy. While TrkB agonists such as 7,8-dihydroxyflavone brought promising results (Jang et al., 2010), systemic administration activates all TrkB signaling pathways and it has been shown increased BDNF-TrkB signaling can lead to seizure activity (Binder et al., 2001). While our approach requires stereotaxic surgery for virus injection, there is extensive research on systemic administration of AAV, which is minimally invasive (Chan et al., 2017). A recent research has also focused on the use of focused ultrasound blood brain barrier opening (FUS-BBBO) for efficient virus delivery. FUS allowed deep penetration in the brain with millimeter spatial precision, and FUS-BBBO combined with systemically administered microbubbles that generate cavitation in blood vessels, can temporarily and reversiblly open BBB (Etame et al., 2012). This allows small molecules such as AAV vector to enter the brain at the specific location where ultrasound is applied. Scablowski et al. has applied this technology to transduce DA neurons in SNC with another chemogenetic probe through I.P injection of AAV in TH-Cre mice(JO Szablowski, 2018). This opens a possibility that virus transduction can be done non-invasively. Current research also focused on designing specific promoters and methods that can target specific neuronal cell types, which does not require a transgenic animal model for cell-type specificity. This approach facilitates translation in a larger animal model or even human patients. Stauffer et al. developed a two viral vectors combination to target DA neurons in non-human primates. The first virus delivered Cre recombinase under the control of a truncated TH promoter and a second vector with a Cre-dependent ChR2 construct (Stauffer et al., 2016).

The use of luminopsins also provided unprecedented advantages in terms of light delivery. Conventional optogenetics utilizes fiber optics and much of light exiting from a single optical fiber is completely attenuated within ∼1 mm from the fiber tip due to light scattering and attenuation that occurs in the brain (Yizhar et al., 2011a), hindering utility of optogenetics in clinical settings. The volumes of SNC in healthy and people with PD are 211 mm^3^ and 180 mm^3^ respectively (Massey et al., 2017), far larger than the volume that can be affected by conventional optogenetics. Although the volume can be improved by enhanced light delivery and red-shifted opsins (Lin et al., 2013), these approaches have only achieved illumination of up to 10 mm^3^ (Galvan et al., 2017). Our previous research showed luminopsin can cover an estimated volume of >20 mm^3^ (Tung et al., 2018). With internally generated bioluminescence, the volume not limited by light scattering or attenuation, but the spread of virally mediated expression.

Translational values of luminopsin has been demonstrated previously. Zenchak et al. showed mice receiving CTZ-activated luminopsin-transduced stem cells in another PD mice model significantly improved in a rotarod test compared to mice receiving vehicle (Zenchak et al., 2018). Similar result has also shown in using luminopsin-transduced stem cells in a stroke model (Yu et al., 2019). These results demonstrated the potential of luminopsin into clinical settings.

While the data presented here is promising, our toxin-mediated PD model does not necessarily represent the neuropathology of PD fully. Even though 6-OHDA model achieves DA depletion, DA neurons loss, and neurobehavioral deficits, it does not affect other regions such as olfactory bulb and locus coeruleus (Surmeier et al., 2017). Furthermore, it fails to exhibit molecular pathology, such as α-synuclein-positive aggregates, a hallmark of PD, and therefore limits the translational value for studying disease-modifying therapies. Recent neuroprotective study shows contradictive results in gene therapy (Decressac et al., 2011; Bartus et al., 2014; Fischer et al., 2017b) using α-synuclein PD model. There are also several clinical trials that used neurotrophic factor treatments, but did not demonstrate a neuroprotective effect or improve symptoms in patients (Thoenen and Sendtner, 2002). This brought attention whether 6-OHDA or other toxin-mediated PD model are useful in assessing neuroprotective treatments(Thoenen and Sendtner, 2002). Even though robust neuroprotection and behavioral improvements were demonstrated with STN-DBS in 6-OHDA rodent model (Fischer et al., 2017a), the treatment failed to protect DA neurons and improve functional deficits in α-synuclein PD model in rodents (Fischer et al., 2017b) while another research with a similar approach showed neuroprotection in α-synuclein PD rodent (Musacchio et al., 2017). While multiple factors can affect these results, future studies will need to include the use of α-synuclein PD model to study whether increasing neuronal activity can diminish neuronal loss due to α-synuclein aggregation.

---

**Supplementary figure 1. SFL expression in HEK293 cells after recombination by Cre**. HEK cells were tranduceed with either pEF1α::Cre-IRES-puroR, pAAV-DIO-SFL, or both of the plasmids and examined under a fluorescent microscope. Expression of the membrane protein, SFL, was observed primarily on the periphery of the cells only when co-tranduceed, confirming specific recombination. As a positive control, HEK cells were also tranduceed with a plasmid with the SFL gene in the normal orientation which does not require recombination.

**Supplementary figure 2**. SFL expression after stereotaxic injection of the AAV vector. DAPI was used for nuclear staining. A) A total of 1.6×10^9^ vg DIO-SFL virs was injected into SNC of TH-Cre mouse. B) A total of 1.6×10^8^ vg DIO-SFL virs was injected into SNC of TH-Cre mouse. A total of 1.6×10^9^ vg DIO-SFL virs was injected into SNC of wild type mouse.

**Supplementary figure 3**. Neuromodulation of SFL-expressing putative dopaminergic cells through bioluminescence. A) Bioluminescence detected through fiber photometry in the SNC. CTZ (20 mg/kg body weight) was injected during the period indicated by yellow rectangle. B) Concomitant increase of firing. Seven putative dopaminergic cells are shown.

