## Supplementary figures for "Chronic activation of dopaminergic neurons via bioluminescence-optogenetics provides neuroprotection in a rodent model of Parkinson’s disease"

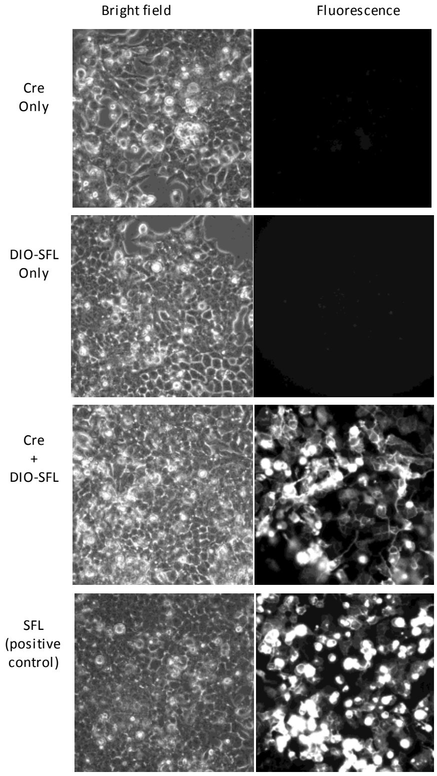


**Supplementary figure 1. SFL expression in HEK293 cells after recombination by Cre.** HEK cells were tranduceed with either pEF1α::Cre-IRES-puroR, pAAV-DIO-SFL, or both of the plasmids and examined under a fluorescent microscope. Expression of the membrane protein, SFL, was observed primarily on the periphery of the cells only when co-tranduceed, confirming specific recombination. As a positive control, HEK cells were also tranduceed with a plasmid with the SFL gene in the normal orientation which does not require recombination.

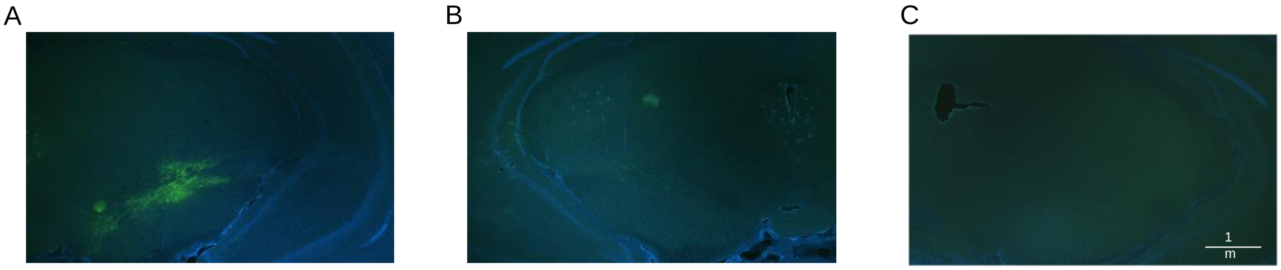


**Supplementary figure 2.** SFL expression after stereotaxic injection of the AAV vector. DAPI was used for nuclear staining. A) A total of $1.6 \times 10^9$ vg DIO-SFL virs was injected into SNC of TH-Cre mouse. B) A total of $1.6 \times 10^8$ vg DIO-SFL virs was injected into SNC of TH-Cre mouse. C) A total of $1.6 \times 10^9$ vg DIO-SFL virs was injected into SNC of wild type mouse.

A

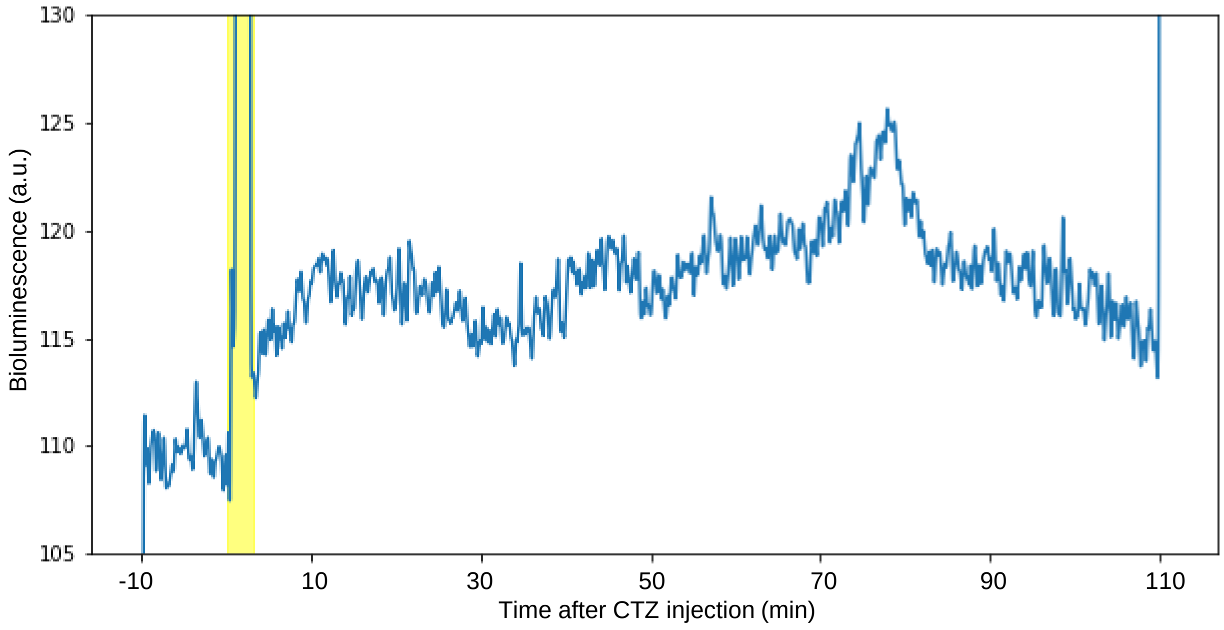


B

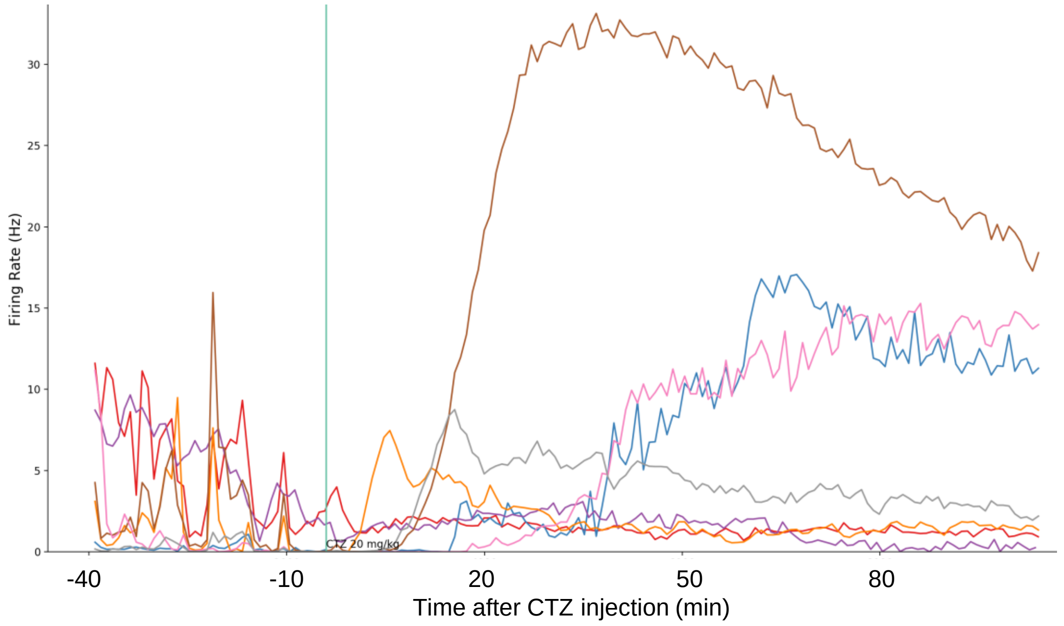


**Supplementary figure 3.** Neuromodulation of SFL-expressing putative dopaminergic cells through bioluminescence. A) Bioluminescence detected through fiber photometry in the SNC. CTZ (20 mg/kg body weight) was injected during the period indicated by yellow rectangle. B) Concomitant increase of firing. Seven putative dopaminergic cells are shown.
